# Enriching stabilizing mutations through automated analysis of molecular dynamics simulations using BoostMut

**DOI:** 10.1101/2025.04.29.651183

**Authors:** Kerlen T. Korbeld, Maximilian J.L.J Fürst

**Affiliations:** Molecular Enzymology, University of Groningen, The Netherlands

## Abstract

Thermostability is a critical goal in protein engineering for applications of biocatalysts and biomedicines. Despite striking advances in biomolecular predictive modeling, reliably identifying stabilizing mutations remains challenging. Previously, molecular dynamics (MD) simulations and visual inspection have been used as secondary filter to improve the success rate of mutations pre-selected by thermostability algorithms. However, this approach suffers from low throughput and subjectivity. Here, we introduce BoostMut (Biophysical Overview of Optimal Stabilizing Mutations), a computational tool that standardizes and automates mutation filtering by analyzing dynamic structural features from MD. BoostMut formalizes the principles guiding manual verification, providing a consistent and reproducible stability assessment. Rigorous benchmarking across multiple datasets showed that integrating BoostMut’s biophysical analysis improves prediction rate regardless of the initial thermostability predictor. Given a modest amount of existing mutant stability data, BoostMut’s performance can be further enhanced with a lightweight machine learning model. Upon experimentally validating BoostMut predictions on the enzyme limonene-epoxide hydrolase, we identified stabilizing mutations previously overlooked by visual inspection, and achieved a higher overall success rate. We foresee BoostMut being used for mutation filtering, as an integrated step in thermostability prediction workflows, and for labelling data to train future predictors.

## Introduction

Proteins are widely utilized in biotechnological and medical applications outside their natural environment, often causing suboptimal performance (Victorino da Silva Amatto et al. 2022). Enzymes, for instance, employed in biocatalysis and synthetic biology, frequently exhibit reduced activity and specificity when acting on non-natural substrates. Another critical limitation in protein applications is instability, causing e.g. low space-time yields in bioconversions or rapid clearance of biological drugs (Woodley 2013; Zaman et al. 2019). Moreover, unstable proteins are poor starting points for enzyme engineering campaigns aimed at enhancing catalytic properties, as such efforts often require the introduction of functionally beneficial, yet destabilizing mutations (Bloom et al. 2006).

A solution to mitigate challenges with instability lies in the use of thermostable proteins, which additionally offer the advantage of being easily enriched or purified through heat denaturation of the expression host’s background proteome (Olichon et al. 2007; Schenkel et al. 2021). While naturally stable homologs from thermophilic organisms are sometimes available, many mesostable homologs display unique functions (Nguyen et al. 2024). Furthermore, extensive prior characterizations of non-stable proteins may discourage switching to new homologs or even *de novo* designed variants. Therefore, engineering existing proteins for enhanced stability is often preferred.

Among methods developed to identify stabilizing mutations, the simplest approach—random mutagenesis followed by screening—primarily yields destabilizing mutations. Although a universal base rate is difficult to determine, estimates based on aggregated stability data suggest that stabilizing mutations (defined as ΔT_m_ > 0 or ΔΔG < 0 kcal/mol) occur at a frequency of 20–30% (Louis and Abriata 2021; Taverna and Goldstein 2002; Tokuriki et al. 2007), although this frequency drops to only 3-5% when neutral mutations (defined as ΔΔG < −1 kcal/mol) are binned separately (Nisthal et al. 2019; Tokuriki and Tawfik 2009). Notably, the practical success rate is lower: for instance, stabilizing the often energetically unfavorable geometries in catalytic sites (Modarres et al. 2016; Shoichet et al. 1995) compromises activity and is thus undesirable. Moreover, because approximately 30% of random mutations impair protein expression by impacting folding or solubility and thus prevent measurement (Velecký et al. 2022), stability datasets likely inflate the apparent proportion of stabilizing mutations.

To improve the identification of stabilizing mutations, numerous computational methods have been developed. Most rely on training or calibrating algorithms on datasets of empirically measured thermostability changes, compiled in various thermostability databases (Bava et al. 2004; Stourac et al. 2020; Xavier et al. 2020). While most energy calculations-based predictors are tuned with these datasets (Guerois et al. 2002; Kellogg et al. 2011; Parthiban et al. 2006), other methods incorporate evolutionary data (Montanucci et al. 2022; Pires et al. 2014). Recent advances in machine learning (ML) have further enabled the integration of structural and evolutionary information from large databases like UniProt and the PDB, rekindling the quest for the long-sought “perfect” stability predictor (Diaz et al. 2024; Jiang et al. 2024; Sun et al. 2023).

Yet, despite the availability of large datasets and continuous algorithmic improvements, the overrepresentation of destabilizing mutations in the data (Caldararu et al. 2020) is the likely cause of a persistent significantly lower accuracy with stabilizing mutations, compared to destabilizing ones (Benevenuta et al. 2023; Diaz et al. 2024). A compounding issue in the field is the use of evaluation metrics such as Spearman correlation and overall accuracy, which fail to account for this imbalance (Broom et al. 2020). The FoldX tool exemplifies this problem: despite strong benchmark performance and widespread use (Buß et al. 2018; Gerasimavicius et al. 2023; Velecký et al. 2024), its predicted stabilizing mutations have an experimental success rate of only ∼29%, while destabilizing mutations are identified correctly ∼69% of the time (Buß et al. 2018). Similarly, the specificity (accuracy for destabilizing mutations) of most other predictors is about twice as high as sensitivity (accuracy for stabilizing mutations) (Khan and Vihinen 2010). Even a state-of-the-art methods large language model predictor only achieved a 44% success rate (45/103) in predicting stabilizing mutations (Li et al. 2010).

In light of these issues, many protein engineering campaigns aim to improve the success rate by combining the primary predictor with a filter step, leveraging the strength of the former in eliminating destabilizing mutations, while applying additional criteria for refinement. In its most simple and common form, a manual, typically visual, inspection assesses mutations before experimental testing, to catch obvious errors. This often overlooked, because understated part of computational engineering pipelines heavily relies on the researcher’s biophysical expertise (Borgo and Havranek 2012; Fischer et al. 2021; Lippow and Tidor 2007). A more advanced filtering mechanism has been proposed in the Framework for Rapid Enzyme Stabilization by Computational Libraries (FRESCO), which uses FoldX and Rosetta to generate a favorable mutation set, followed by high-throughput molecular dynamics (MD) simulations and visual inspection to incorporate structural dynamics (Wijma et al. 2014; Wijma et al. 2018). This method has consistently led to substantial thermostability gains, with one campaign producing a 10-fold mutant exhibiting a ΔT_m_ of +51 °C (Aalbers et al. 2020).

However, although visual inspection improves thermostability predictors’ success rates, it remains limited by subjectivity, labor intensity, and low throughput. The reliance on human judgment constrains speed, scale, and reproducibility, making systematic high-throughput screening impractical. To overcome this issue, we developed BoostMut (Biophysical Overview of Optimal Stabilizing Mutations), a secondary filter that analyzes structural features of MD simulated mutations. BoostMut formalizes the inspection principles, providing a consistent and reproducible method for evaluating mutations generated by primary predictors. BoostMut outputs a set of interpretable biophysical metrics that can rationalize the effect of mutations and serve as structured input to train ML models. We evaluated BoostMut on 1,584 mutations across six proteins and established that it outperforms visual inspection and commonly used predictor scores in ranking. We demonstrate its effectiveness by experimentally analyzing 18 previously untested mutations in limonene epoxide hydrolase, and show that BoostMut filtering selects stabilizing mutations with a success rate of 50% in this protein.

## Results

At its core, BoostMut analyses various biophysical properties of protein structure ensembles such as MD trajectories, to compare the stability of a set of mutants to the wild type. While running even short MDs for all possible single mutants would typically be too computationally expensive, it becomes feasible once a narrower set of mutations has already been pre-selected. MDs and BoostMut are therefore specifically meant as a filter after already having selected mutations by a given primary predictor (Figure 1a).

**Figure 1.**
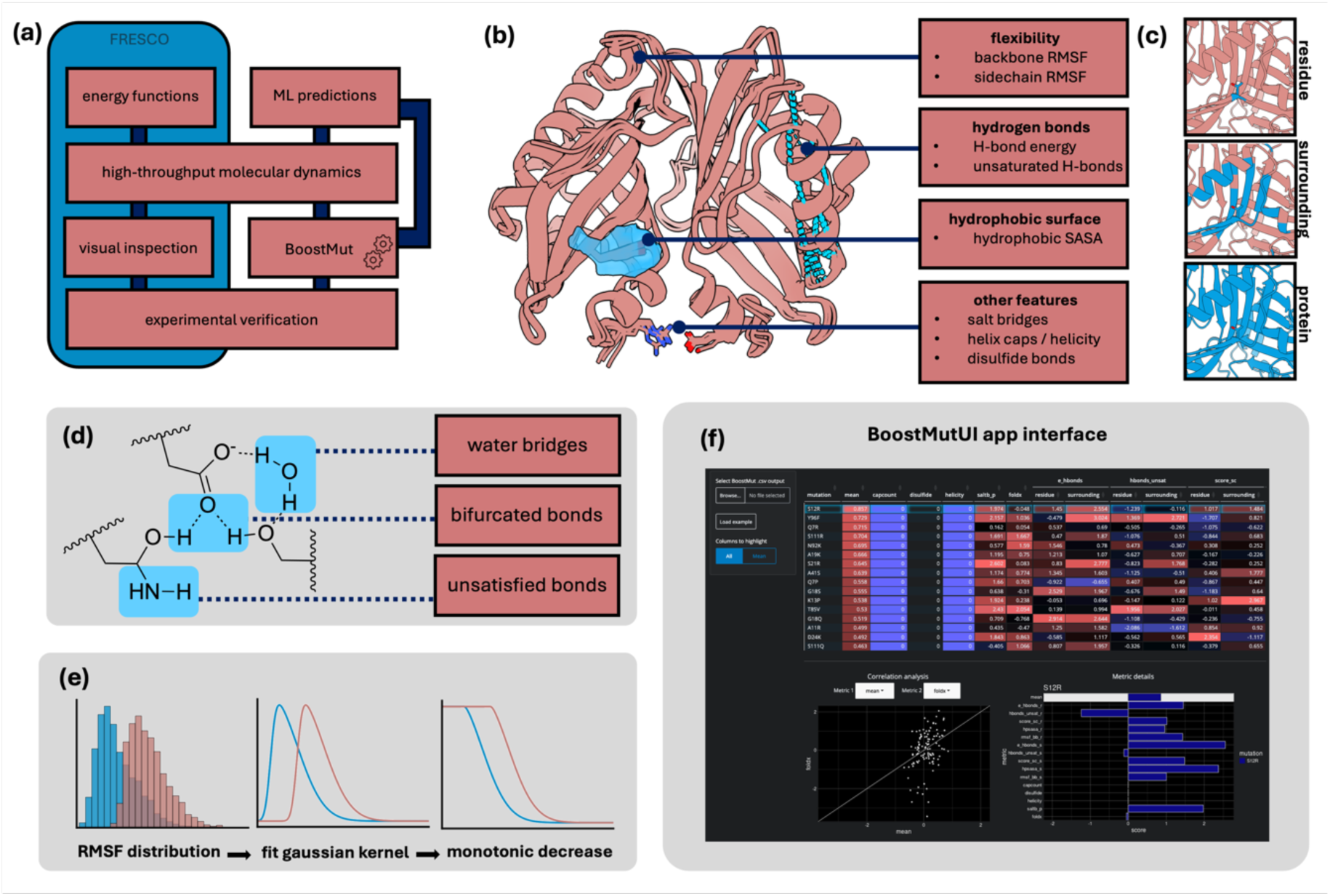
(a) Overview of the FRESCO and BoostMut pipelines. (b) The BoostMut metrics. (c) Metrics are evaluated at the residue, 8 Å surrounding, or the entire protein level. (d) Evaluation of protein hydrogen bonding distinguishes water-mediated H bonds, bifurcated H bonds, and unsaturated donor/acceptors. (e) To obtain benchmark curves, the distribution of the property is fitted to a gaussian kernel, and a monotonic decrease is guaranteed. (f) BoostMut’s online user interface visualizing the output.

The metrics analyzed by BoostMut are based on physical properties of proteins known to affect thermostability and widely used as screening criteria when assessing mutations (Floor et al. 2014; Wijma et al. 2018). They mainly comprise improvement of the hydrogen bond (H bond) network via increased intramolecular or reduced unsatisfied donor/acceptor groups, prevention of increased protein flexibility, and minimization of solvent-exposed hydrophobic residues (Figure 1b). BoostMut encapsulates these and other criteria into a formalized set of metrics, whose output can be used for scoring and ranking, or as input for ML methods.

We implemented our analyses via the MDAnalysis python library (Gowers et al. 2016), which provides a framework for analyzing MD trajectories and allows BoostMut to work across different topology and trajectory formats. For each metric, the difference between the mutant and the wild type, averaged over each frame of the trajectory, is reported. BoostMut always calculates at three distinct levels of granularity: the mutated residue itself, its local environment, or taking the entire protein into account. For each selection, a more complete effect of the mutation on its surrounding is obtained, but also the amount of noise increases (Figure 1c).

The first metric we assess is hydrogen bonding, which is known to increase protein stability by decreasing the enthalpy of the folded state (Pace et al. 2014). Yet, these contributions are often small (Hubbard and Kamran Haider 2010), due to the desolvation penalty for disrupting water bonding in the unfolded state (Bolen and Rose 2008). For the same reason, unsatisfied H bonds favor the unfolded state (McDonald and Thornton 1994). We therefore assess H bond changes in two metrics: the increase in internal protein-protein bonds and the decrease of unsatisfied H bonds. BoostMut estimates intramolecular binding energy by assessing angles and distances, assuming 25 kJ/mol for ideal interactions and scaling down based on deviations (Fleming and Rose 2005; Hubbard and Kamran Haider 2010). Further refinements account for the effect of bifurcated H bonds (Feldblum and Arkin 2014) and inclusion of the binding energy of water-mediated H bonds (Petukhov et al. 1999) (Figure 1d). Unsatisfied H bonds are defined as all potential donor and acceptors (N, O, S, and their bound H atoms within the protein) without interactions.

Protein flexibility is another key stability marker, as structural fluctuations can expose the hydrophobic core and cause unfolding (Bommarius and Paye 2013). Increased flexibility typically correlates negatively with stability (Kamerzell and Russell Middaugh 2008; Karshikoff et al. 2015; Yu and Huang 2014), an insight leveraged in previous thermostability designs (Yu and Huang 2014). Accordingly, we use lower root mean square fluctuations (RMSF) of the protein backbone Cα atoms during the MD as an indicator of stabilization. Separately, we examine sidechain flexibility, where an increase may indicate loss of favorable local interactions. To compare flexibility between different amino acids, we normalized RMSFs using a custom benchmark dataset of expected flexibility per amino acid and degree of surface exposure (Figure 1d). This benchmark set was generated by running MDs on a curated set of protein structures, obtained by filtering the protein data bank (PDB) for protein-only structures with high resolution (<=1 Å) and a minimum size (>100 aa, to retain a well-defined hydrophobic core). With this set of 77 proteins, we generated the distributions of expected RMSFs for each amino acid type binned into “exposed”, “partially buried”, or “fully buried” based on a relative solvent-accessible surface area (SASA) of >20% of the total surface, 0-20%, or 0%, respectively (Chen and Zhou 2005; Zhang et al. 2009). To each of the resulting 60 benchmark distributions, we fitted a gaussian kernel, scaled to values between 0 and 1, and enforced a monotonically decreasing curve (Figure 1e, Supplementary Figure S2), to only penalize unusual flexibility, but not rigidity.

BoostMut further assesses hydrophobic exposure. As the main driver of protein folding, hydrophobic interactions promote core packing, while polar residues remain solvent-exposed. Consequently, exposed hydrophobic residues often destabilize proteins (Qing et al. 2022). Although some studies report a prevalence of mutations introducing hydrophobic surface residues (Broom et al. 2020; Nisthal et al. 2019), hydrophobic exposure is generally undesirable due to reduced solubility and aggregation (Broom et al. 2017). BoostMut therefore penalizes increased hydrophobic exposure by summing the SASA of all C atoms and all H atoms bound to C, relative to values again obtained via a custom benchmark set, which compensates for polarization and other effects ignored in this simplified definition (Li et al. 2010). For the benchmark, we generated normalized and monotonically decreasing distributions, reflecting the typical surface exposure of each amino acid (Supplementary Figure S3).

Besides these main metrics, BoostMut includes additional minor checks. Since α-helices rely on local interactions, the sequence-structure relationship manifests stronger than for other secondary structures (Ismi et al. 2022). We implement this knowledge by assessing helix propensity—a scale assigning an energetic penalty to specific amino acids introduced into α-helices (Pace and Scholtz 1998). While most residues exhibit mild penalties (0–4 kJ/mol), proline is a major exception (13.22 kJ/mol), allowing BoostMut to prevent proline-induced helix breakage (Woolfson and Williams 1990). Another determinant of helix stability are helix-capping motifs, which complete a helix termini’s H-bonding network, and whose disruption introduces destabilizing unsatisfied H bonds (Aurora and Rosee 1998). As these effects may only emerge in extended simulations, BoostMut explicitly evaluates whether a mutation interferes with helix capping. Helix propensity is only checked for residues at least five positions from helix termini, ensuring the evaluation to occur independently from helix caps.

In addition to H bonds, proteins are also stabilized via ionic interactions between positively and negatively charged residues (Kumar and Nussinov 1999; Panja et al. 2020). To prevent mutations from disrupting these salt bridges, BoostMut tracks their average number throughout the MD. As is common, we define a salt bridge as any occurrence of a negatively charged oxygen atom in aspartic or glutamic acids closer than 4 Å to a positively charged amine groups in arginines, lysines, or histidines (Barlow and Thornton 1983; Makhatadze et al. 2003).

We also consider disulfide bonds, which stabilize proteins via covalently linked cysteine residues, thus favoring the folded conformation by reducing unfolded-state entropy (Betz 1993; Dombkowski et al. 2014; Feige et al. 2018). While typically designed by specialized algorithms rather than general stability predictors (Dombkowski et al. 2014; Wijma et al. 2014), we implemented an identification step in the input structure to prevent their disruption.

BoostMut evaluates each of these metrics for the three selections, except for non-fluctuating metrics (helix analysis and disulfide bonds), which are only returned for the entire protein context. All metrics are then reported as the difference between mutant and wild type, returning 0 for a mutation with neither detrimental nor beneficial effects. While the individual outputs may inform users on specific effects of a given mutation, the main output of BoostMut is a composite score for ranking mutations. To that end, each metric is normalized by dividing by the standard deviation within the set, yielding z-score-like values with the wild-type score (0) as the reference. The sign is adjusted such that a positive score consistently indicates a stabilizing effect. If the primary stability predictor also outputs a numerical score, it may be included as an additional metric for the ranking after normalization. The final BoostMut score is simply the mean of all metrics and is used to rank the mutations.

A BoostMut-containing pipeline then runs as follows: 1) a primary predictor selects an initial set of mutations via classification or based on a cutoff, 2) each selected mutant is simulated in (typically short) MD simulations, with length and number of replicates based on available resources, followe by 3) BoostMut scoring of the mutations, enabling users to select any number of top-ranked mutations for experimental validation. The required benchmark data can be obtained from our implementation for trajectories up to 5x 1 ns. We distribute BoostMut as a pip-installable python package with a command line interface allowing customization of analyses, selections, and trajectory lengths via command-line flags. The main output is a tabular file containing raw metric scores and composite rankings. We also developed a simple app hosted on https://fuerstlab.shinyapps.io/BoostMutUI/ for interactive exploration of the results (Figure 1f)

To assess BoostMut’s effectiveness and establish optimal parameters, we evaluated its performance using data we had available from previous thermostability engineering projects applying the FRESCO workflow. As this protocol employs energy calculations, short MD simulations on mutants passing a ⎼5 kJ/mol energy cutoff, and visual inspection ensuring structural feasibility and rigidity, the experimentally tested mutations from this set are highly pre-selected. We obtained data from FRESCO campaigns that stabilized three biotechnologically relevant enzymes: limonene-1,2-epoxide hydrolase (LEH) (Wijma et al. 2018), alcohol dehydrogenase 99 (ADH) (Aalbers et al. 2020), and 5-hydroxymethylfurfural oxidase (HMFO) (Martin et al. 2018), resulting in 1251 threshold-passing point mutations, of which a total of 339 mutations were experimentally verified.

While the original pipeline producing this dataset simulated 5 replicates of only 50 ps, we were curious to examine the impact of MD simulation length on prediction accuracy, and re-simulated the 106 threshold-passing LEH mutations for 5x 1 ns each.

To track convergence of the ranking, we analysed the Pearson correlation coefficient (PCC) between variable trajectory lengths and the final 1 ns trajectory (Figure 2a). Noticing that the whole-protein selection converges significantly more delayed (Figure 2a), we chose to exclude it in BoostMut by default. An exception is made for salt bridges, which are less subject to noise due to their relative rarity, but are frequently absent at the residue or local level. Nevertheless, some mutations showed a clear score change at longer simulation lengths, with the strongly stabilizing N92K mutation (ΔT_m_ +7.3°C) becoming a pronounced outlier at longer time scales (Figure 2b). The higher score was primarily caused by this mutant’s lysine sidechain adopting a distinct conformation that cemented subunit interaction, which led to a significantly reduced local flexibility as the simulation progressed (Supplementary Video 1).

**Figure 2.**
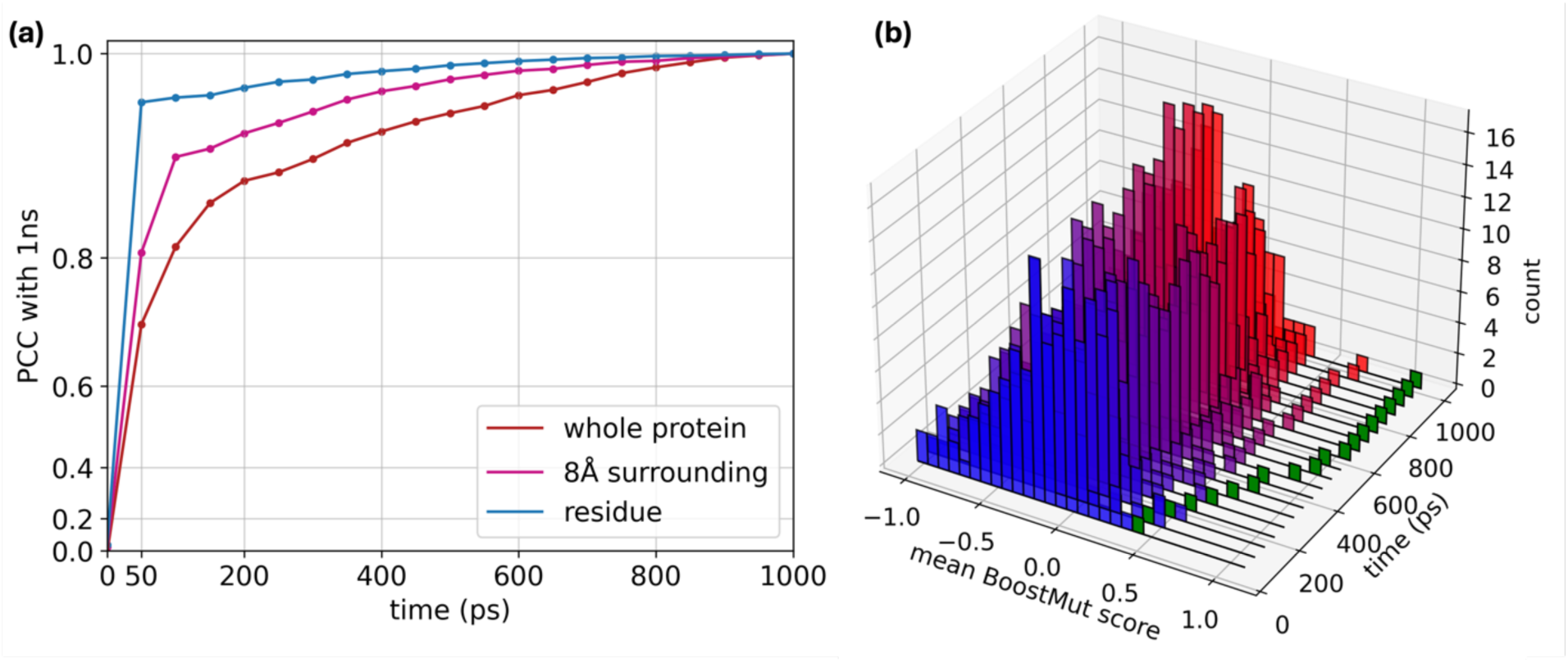
(a) Pearson correlation coefficient (PCC) tracking the similarity of BoostMut ranking on the LEH dataset to the final ranking at 1000 ps for each MD length, calculated for the three selections in intervals of 50 ps. Significantly higher PCCs for the residue and surrounding selections indicate faster convergence of the outcome compared to the whole protein selection. (b) Distribution of BoostMut scores for differing simulation times. The highly stabilizing mutation N92K (shown in green) is an example of a mutation benefitting from longer simulations times.

**Figure 3.**
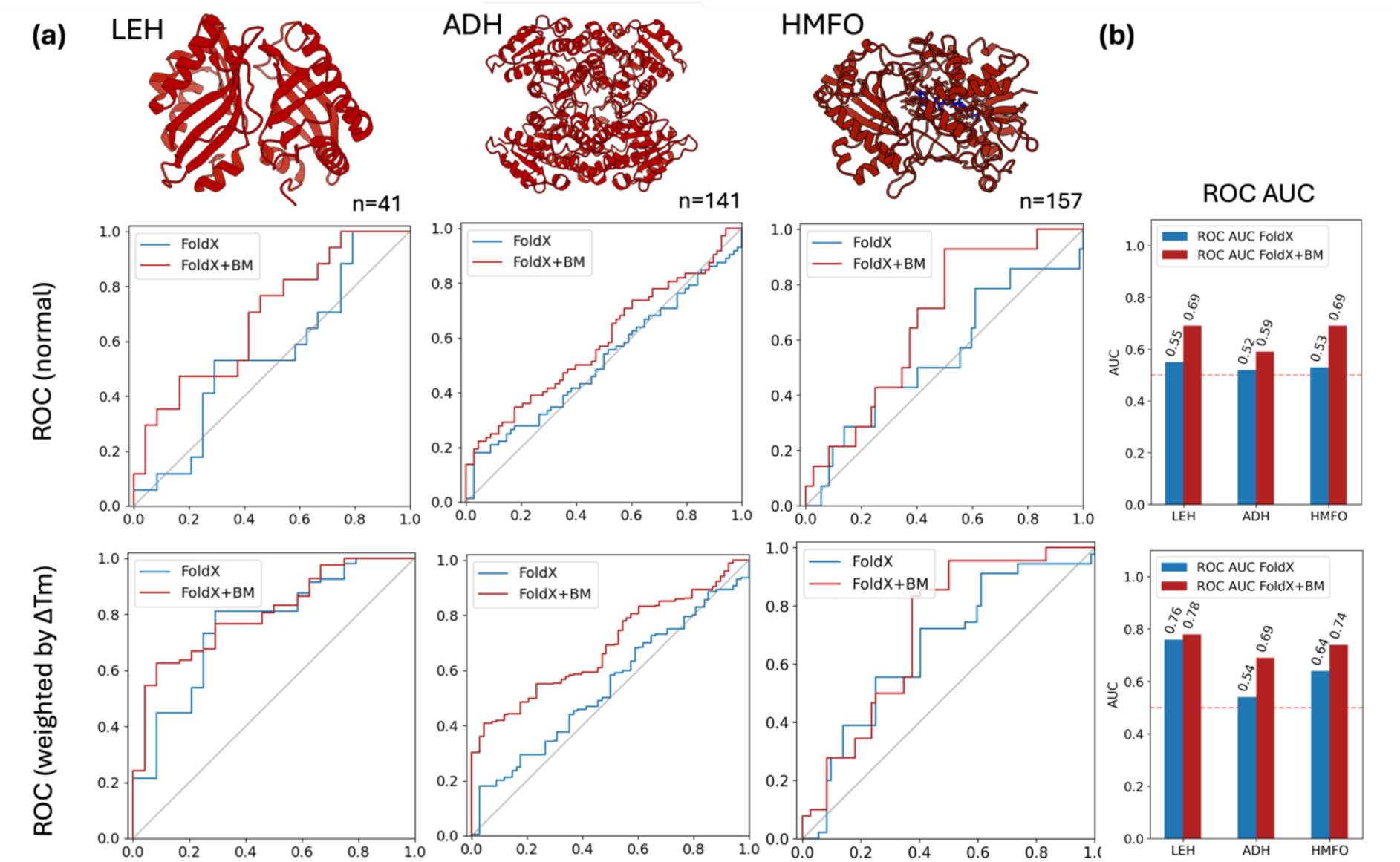
(a) Normal and weighted ROC curves for three proteins from previous stability engineering campaigns (LEH, ADH, HMFO) upon re-ranking by BoostMut (BM). The weighted ROC curves scale each mutation by their respective ΔT_m_. (b) The improvements upon reranking by BoostMut result in a consistent increase in AUC. The increase in AUC from the normal to weighted ROC curves indicates a preferential sorting of highly stabilizing mutations.

We tested BoostMut on all 1251 mutations in our dataset, using data and benchmarks adjusted to the original protocol of 5x 50 ps per mutation (0.3 µs total simulation time). While the MDs correspond to a computational budget of a few days on a high-end consumer GPU (Abraham et al. 2015; Eastman et al. 2017), BoostMut analysis of the combined 125,100 snapshots of all the mutations required approximately 60 total CPU hours, or ∼3 minutes per mutation per CPU. The original visual inspections selected 339 mutations for experimental validation, forming our benchmark set. To avoid overestimating predictor performance, we did not use metrics like Pearson correlation, classification accuracy, or error rates (Diaz et al. 2024), which may let predictors that simply frequently classify mutations as destabilizing appear accurate. Instead, we used receiver operating characteristic (ROC) curves, which are independent of class distribution. The area under the ROC curve (AUC) quantifies ranking ability: 1 indicates perfect classification, 0.5 random sorting, and values below 0.5 reflect a preference for destabilizing mutations. To be deliberately conservative, we treated neutral mutations as unsuccessful, defining the classes as stabilizing and not stabilizing. As thermostability is a quantitative metric, we additionally propose a new metric, stability-weighted ROC curves, where stabilizing mutations are scaled by their ΔT_m_ or ΔΔG contribution. This approach rewards predictors that prioritize highly stabilizing mutations with higher AUCs, offering a metric of practical benefit in real-world thermostability engineering.

We first generated standard ROC curves for the FoldX scores of the 339 experimentally tested mutations. As expected, AUC values were low, only slightly above 0.5 (Figure 4). This is unsurprising, as all mutations had not only been visually approved, but also already passed a ΔΔG cutoff of –5 kJ/mol—meaning FoldX already classified them as stabilizing, and the scores correlate poorly within this narrow, pre-selected range. This outcome thus reflects a broader limitation: predictors trained on datasets dominated by destabilizing mutations tend to perform poorly when ranking stabilizing ones. We then probed the outcome upon BoostMut filtering and found elevated AUC scores for all three proteins, increasing the average (sample-size adjusted) AUC score from 0.53 to 0.65 and 0.61 to 0.72 for unweighted and weighted ROC curves respectively. Notably, the stability-weighted curves displayed higher AUCs regardless of ranking method (indicating that both FoldX and BoostMut prioritize highly over modestly stabilizing mutations), but BoostMut consistently ranked stabilizing mutations more effectively than FoldX alone. These results imply that BoostMut can increase both quantity and quality of stabilizing hits in a highly pre-selected set, where primary predictors struggle to accurately distinguish mutation effects.

**Figure 4.**
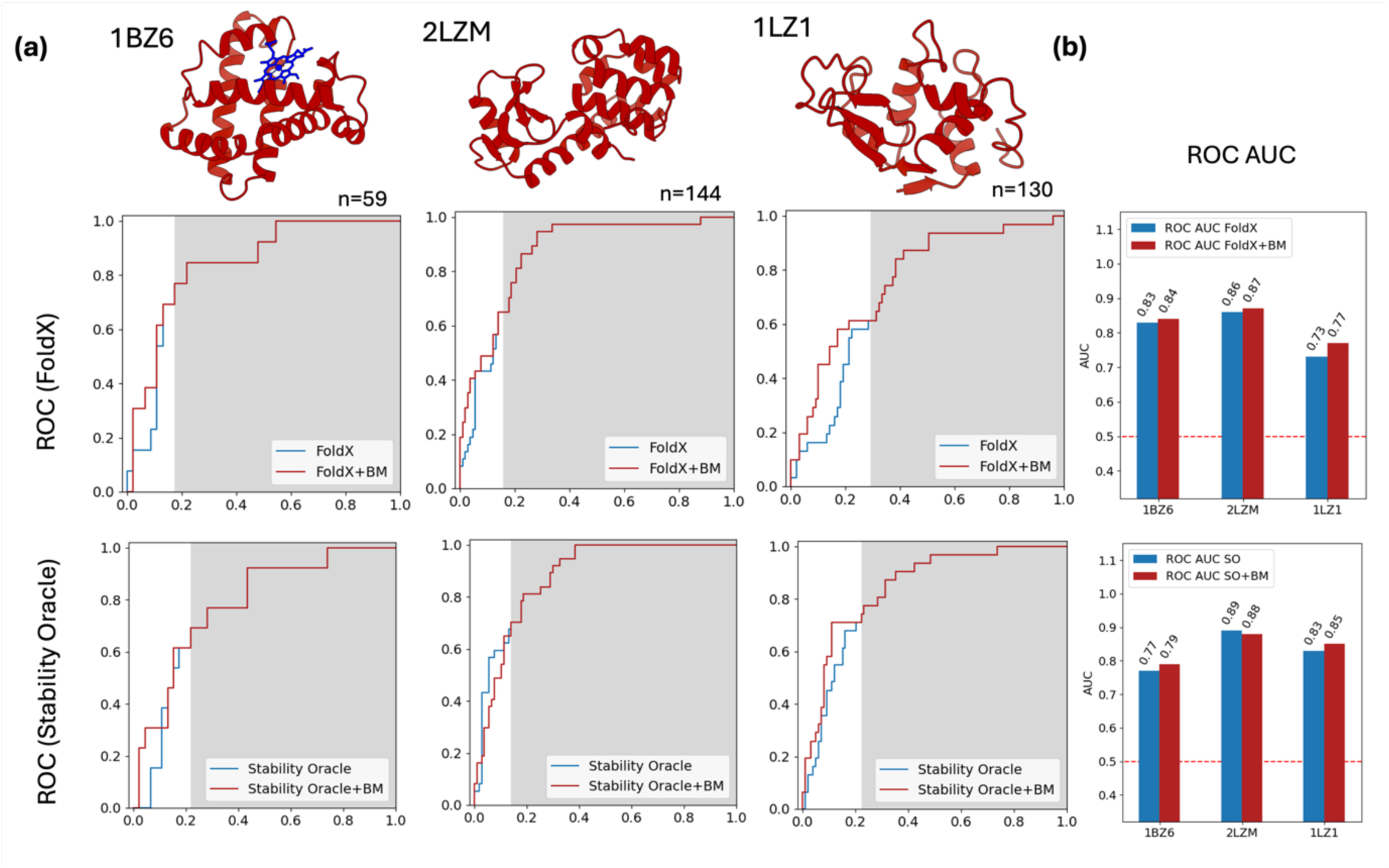
(a) ROC curves for three proteins from the T2837 test set, using either FoldX (top) or Stability Oracle (bottom) as the primary predictor. All mutations with a predicted ΔΔG < 0 were scored with BoostMut (white area), otherwise by the primary predictor (grey). (b) For 5/6 cases, the addition of BoostMut resulted in a small increase in ROC AUC.

Since BoostMut functions as secondary filter and requires MDs, we could not directly compare it to data from online databases (Diaz et al. 2024; Velecký et al. 2024). To expand the benchmark nonetheless, we evaluated the performance of BoostMut on a subset of the T2837 dataset used to benchmark Stability Oracle, a recent state-of-the-art ML method (Diaz et al. 2024). This data comprises mutations with both positive and negative FoldX scores, allowing us to generate ROC curves that represent the combined effect of a complete BoostMut-incorporating pipeline in which it ranks a primary algorithm’s selection. We selected three proteins with over 100 mutations (PDB IDs 1BZ6, 1LZ1, 2LZM) and excluded those near cofactors, yielding 333 mutations. Since only few passed the default thresholds, we used a relaxed cutoff of 0 kJ/mol, resulting in 107 pre-selected mutations for FoldX and 102 for Stability Oracle. After recreating these mutant’s structures with FoldX and running 5x 50ps MD simulations, we ran BoostMut to rank them.

In this analysis, the ROC curves led to significantly higher AUCs than for the FRESCO data, as one would expect with a significantly lower degree of preselection, which creates a more mixed set of mutations (Figure 4). In line with the intended BoostMut use, we reranked only the cutoff-passing mutations (Figure 4, white area), while the “rejected” mutations (Figure 4, grey area) kept their original order. Here, the weighted ROC curves (Figure S1) showed a less pronounced difference, reflecting this set’s narrower distribution of determined ΔΔG values. In agreement with what was reported, we find Stability Oracle to slightly outperform FoldX on the whole: the average (sample-size adjusted) AUCs were 0.84 and 0.80 for Stability Oracle and FoldX, respectively. Notably, the additional re-ranking by BoostMut resulted in a small but consistent improvement, raising the average AUC from 0.80 to 0.82 for FoldX and raising the average AUC from 0.84 to 0.85. Thus, BoostMut is apparently able to enhance the performance of diverse primary predictors and across different levels of preselection.

Since one of our key goals was replacing visual inspection for full automation in engineering pipelines, we directly compared BoostMut’s performance to human filtering. Thus, we scored all 1251 MD-simulated FRESCO mutations with BoostMut and selected the same number of top-ranked mutations as chosen manually: 41 for LEH, 141 for ADH, and 157 for HFMO. Notably, fewer than half of the two sets’ mutations overlapped (Figure 5A). Yet, within the mutations selected by BoostMut with known ΔT_m_, stabilizing mutations were significantly enriched: 10/20 for LEH, 36/66 for ADH and 4/27 for HMFO, at an average success rate of 50/103 or 49% (Figure 5B). However, as 225 of the BoostMut-selected mutations and 912 of the remaining 1251 MD-simulated FRESCO mutations lack experimental data, the true success rate remains unclear. Consequently, we selected a feasible number of top-ranked, uncharacterized mutations from LEH for experimental verification, where the smaller initial dataset would cause the new mutations to constitute a larger proportion of the whole. Among the 19 so-selected mutations, Q7R, the highest-ranked untested BoostMut mutation, had already been reported stabilizing by Sun et al., and we thus included its ΔT_m_ (Sun et al. 2023).

**Figure 5.**
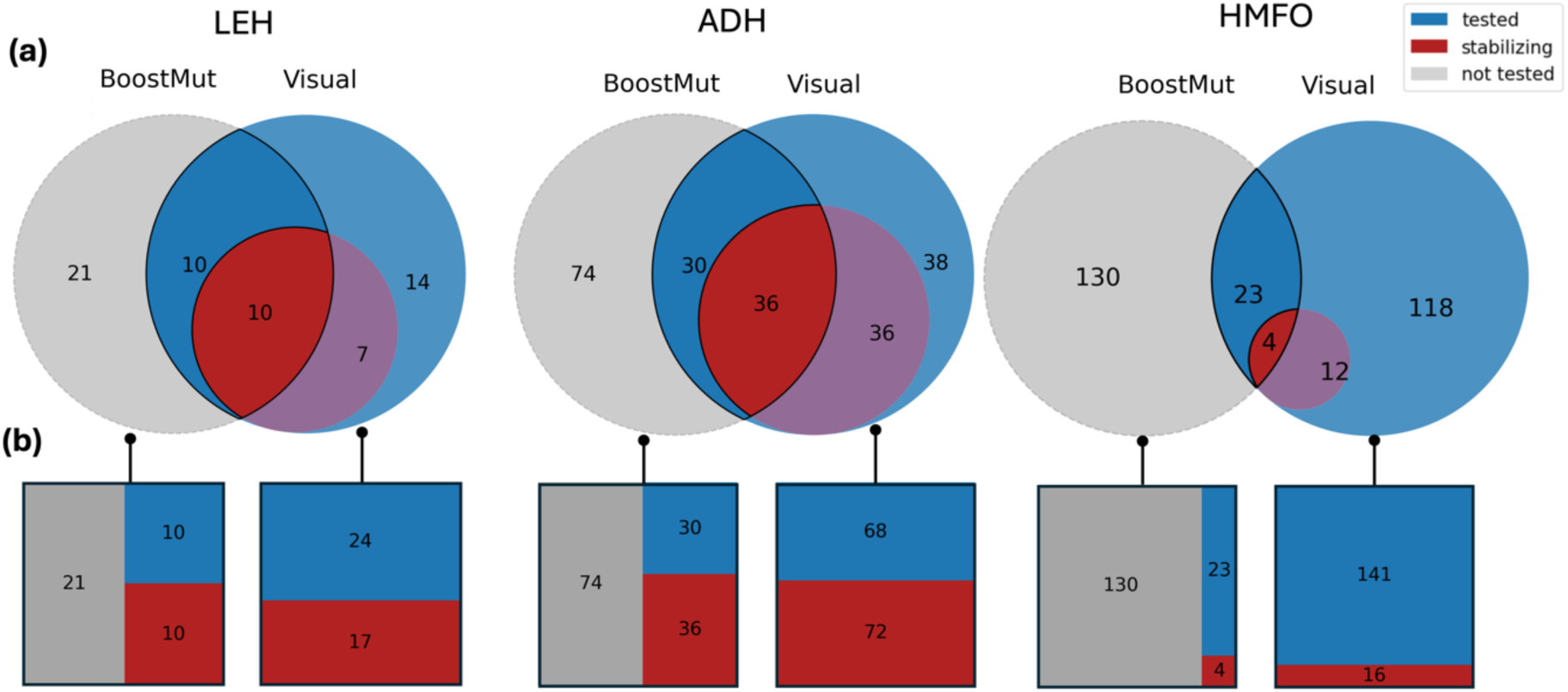
(a) Limited overlap in the mutations selected by visual inspection and BoostMut for three proteins from previous FRESCO campaigns (LEH, ADH, HMFO). (b) When comparing the mutations for which the experimental outcome is known, the fraction of stabilizing mutations is higher for BoostMut.

We generated *E. coli* expression constructs for all new mutations via QuikChange of a pBAD expression vector for LEH, following established procedures (Wijma et al. 2014; Wijma et al. 2018). All mutants were expressed with a 6xHis-tag, purified by affinity chromatography, and desalted. Apparent melting temperatures were measured via a thermofluor assay (Pantoliano et al. 2001). Of the 19 newly selected mutations, 7 reduced T_m_, 4 were neutral, and 8 were stabilizing (Figure 6a), including 2 with ΔT_m_ increases >4 °C—among the most stabilizing ever reported for LEH. Three mutations occurred at a previously identified stabilizing site, while five were at new positions (Figure 6b). Notably, the top four BoostMut-ranked mutations were all stabilizing. Upon including the new mutations into the overall comparison, BoostMut’s success rate for LEH increased further, with 20/40 (50%) stabilizing, compared to 19/46 (41%) for visual inspection (Figure 6c).

**Figure 6.**
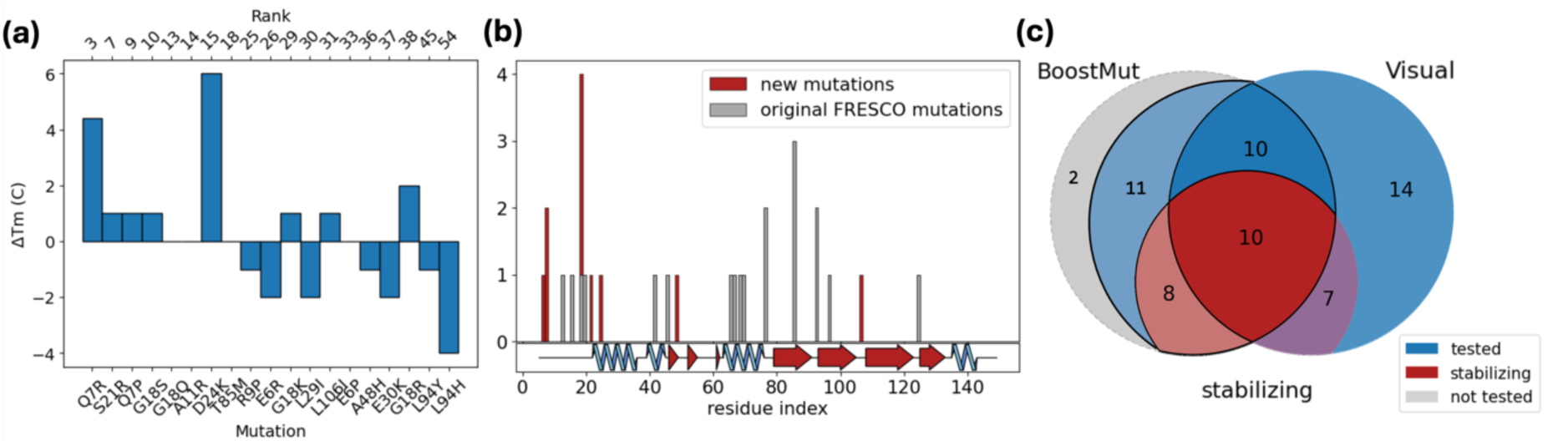
(a) ΔT_m_ of 19 the top ranked BoostMut mutations for LEH. (b) location of all stabilizing mutations on the protein. Previously known mutations are grey, new ones red. (c) Venn diagram of the overlap between the mutations selected by visual inspection and by BoostMut when including the experimental data from the new mutations.

For a more thorough comparison, we again resorted to ROC analysis. However, despite the additional data, ROC analysis with all 108 cutoff-passing LEH mutations remained complicated by the remaining 58/108 variants not selected for experimental verification by either method. To estimate possible outcomes, we programmatically evaluated all combinatorial scenarios, thus establishing upper and lower AUC bounds (Figure 7a). Since the original visual inspection followed a binary accept/reject approach, we ranked the mutations by FoldX scores, after placing all accepted before the rejected mutations. To compare more widely, we also evaluated this dataset with other structure-based stability predictors. We thus applied Pythia (Sun et al. 2023), DDGun (Montanucci et al. 2022), DDMut (Zhou et al. 2023), DUET (Pires et al. 2014), Mupro (Cheng et al. 2006), mCSM, and PremPS (Chen et al. 2021) (Figure 7b) as an alternative means to score the mutations. Besides FoldX and Pythia, the BenchStab package was used to run the predictors (Velecký et al. 2024).

**Figure 7.**
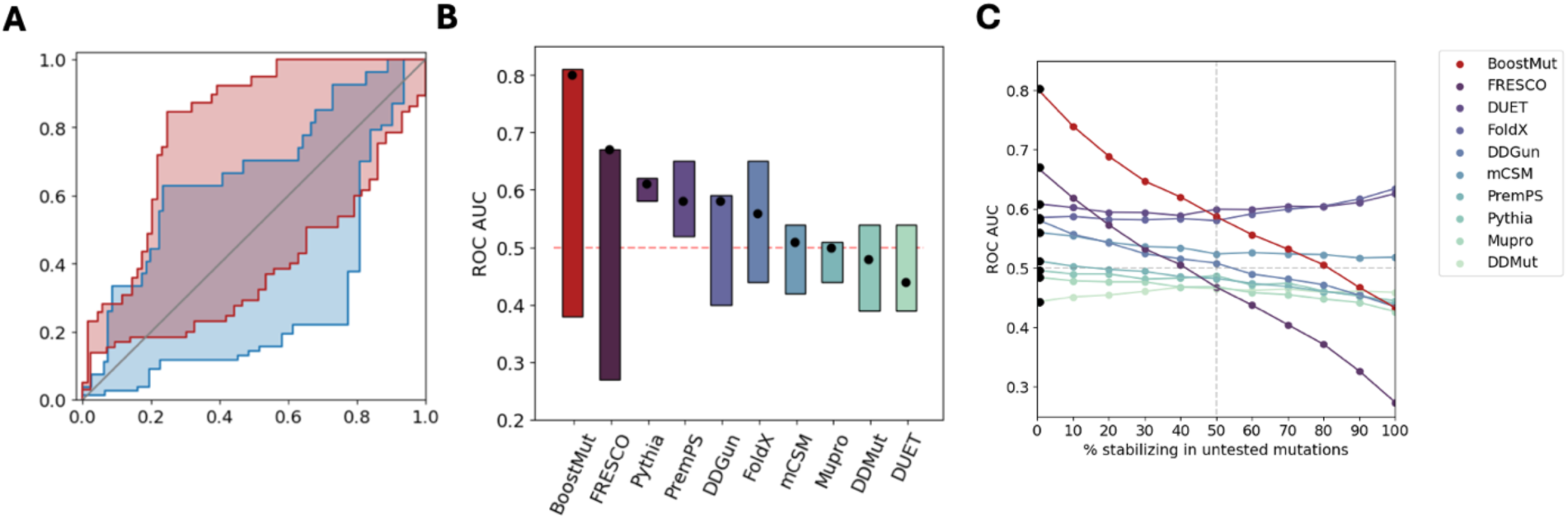
(a) potential ROC curve ranges of visual inspection (blue) vs. BoostMut (red), depicting the best and worst-case scenarios for the remaining untested mutations. (b) Potential ranges of ROC AUC values for BoostMut, visual inspection, and other primary predictors. The scenario where all the remaining untested mutations are destabilizing is shown as black dots. (c) Changes in ROC AUC if an increasing fraction of untested mutations were actually stabilizing. BoostMut outcompetes all other methods until the scenario where 50% of untested mutations are stabilizing.

Our initial goal was to match human inspection performance, but BoostMut ultimately outperformed it. Across all possible outcomes for the untested mutations with unknown ΔT_m_, BoostMut scoring consistently resulted in higher AUC values than visual inspection, with bounds of 0.80–0.43 compared to 0.67–0.27, respectively (Figure 7). Regardless of whether the untested mutations would turn out to be all stabilizing or destabilizing, BoostMut retains an approximately 0.12 AUC advantage (Figure 7c). This indicates that BoostMut would likely outperform human judgement regardless of how many of the untested mutations would be identified as stabilizing. The large span in possible AUC values is caused by the low ranking of the remaining mutations rejected by both BoostMut and visual inspection, since introducing new true positives near the end of the ROC curve has a larger detrimental effect on AUC. On the contrary, the other predictors are less sensitive to these fluctuations (Figure 7b), reflecting the fundamentally different algorithmic determination. However, we also found that all tested tools suffer from the same weakness in accurately predicting stabilizing compared to destabilizing mutations in this pre-selected set, reinforcing the utility of secondary filtering. We find that as long as fewer than 50% of untested mutations are stabilizing, BoostMut remains the best choice (Figure 7c). Given that all untested mutations were rejected by visual inspection, their expected stabilizing fraction should be at most 30%, based on the overall average success rate when using the default –5 kJ/mol FoldX cutoff. However, if visual inspection actually served its intended purpose and led to an enrichment, this fraction would be even lower. If more than 50% of the remaining mutations were stabilizing, Pythia and PremPS—the two best-performing predictors in our analysis—would outperform BoostMut. However, for this to occur, visual inspection would have needed to reject stabilizing mutations at a high rate, effectively lowering rather than enriching the success rate. Since this is unlikely, we expect the true stabilizing fraction to fall between 0–30%, rendering BoostMut as best choice for thermostable mutant filtering.

To better understand BoostMut’s strengths and limitations, we examined illustrative examples of its performance (Figure 8). A notable success is L150F in ADH, which increased stability by 14.2 °C. BoostMut scored it highly across several metrics, particularly for reduced hydrophobic exposure and lower backbone RMSF—consistent with visual inspection notes describing improved hydrophobic packing as the phenylalanine fills a cavity. S21R is a by BoostMut newly found mutations in LEH (+1 °C ΔT_m_) that was manually rejected (Figure 6a). The main contributions to its positive BoostMut score come from the salt bridge and the H bond energy scores. While this improvement was also identified in the visual inspection, the mutation was rejected for a perceived increase in flexibility. However, both the backbone RMSF and the sidechain BoostMut scores are favorable, demonstrating the value of a more objective metric in evaluating flexibility. Other examples also highlight limitations of MD-based screening. For instance, G463E in HMFO was favored by both visual inspection and BoostMut due to apparent hydrogen bonding and salt bridge formation, but the mutation was experimentally found to decrease stability. To systematically compare BoostMut and visual inspection, we manually examined all 43 mutations in LEH where their selections differed. In most cases (29/43), the underlying observations were largely in agreement, but differences in prioritization led to divergent outcomes. The remaining 14 cases involved direct conflicts, primarily in flexibility or hydrogen bonding, both of which are challenging to assess visually.

**Figure 8.**
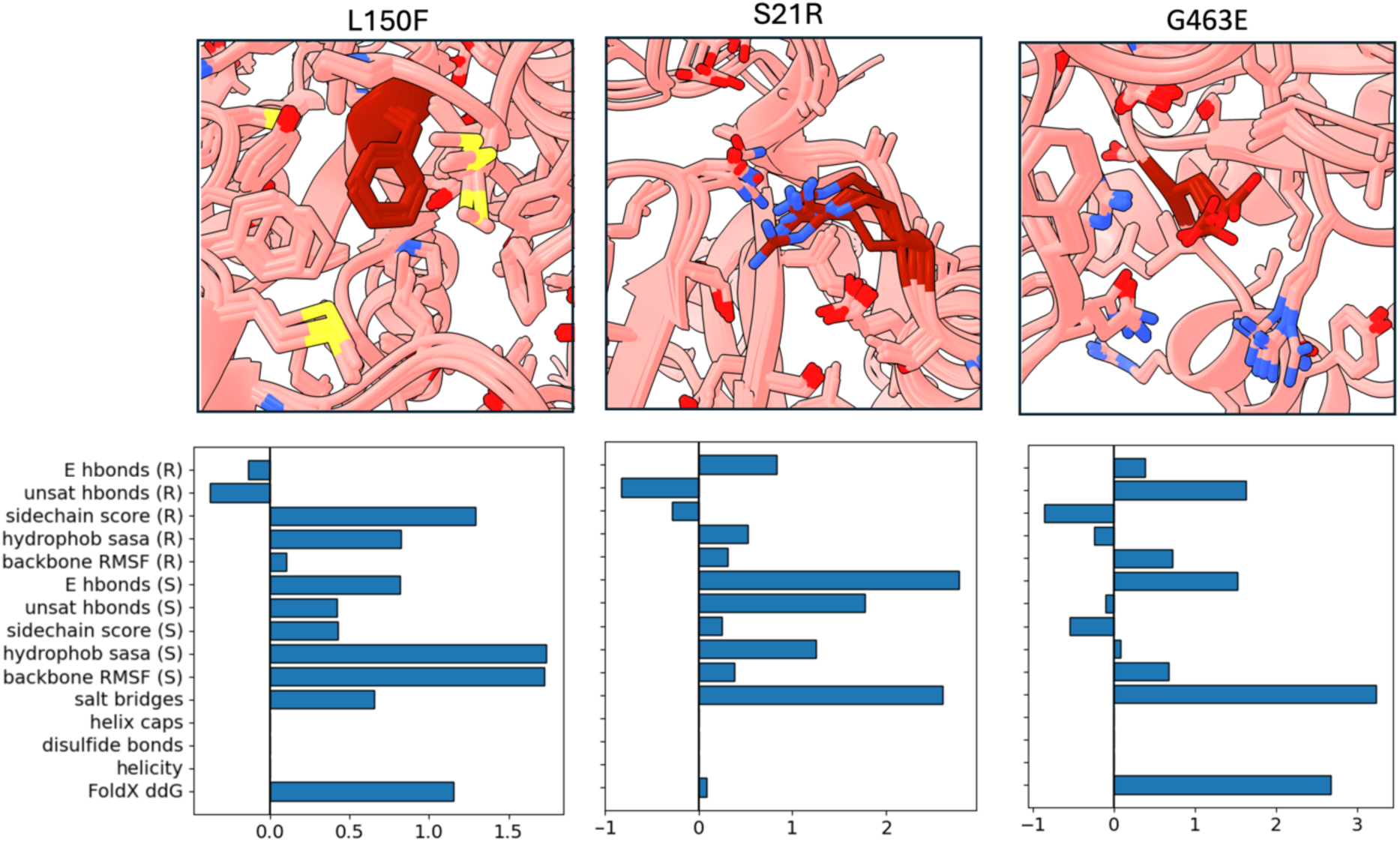
The scaled metrics for 3 selected mutations. L150F and S21R are both stabilizing mutations, successfully identified by BoostMut. G463E is a destabilizing mutation incorrectly classified as stabilizing by both visual inspection and BoostMut.

We reasoned that BoostMut’s consistent assessment of biophysical stabilization mechanisms may provide a useful dataset for machine learning. While BoostMut by default simply averages all metrics to a final score, we asked whether we can train a model to optimize metric weights. Although fully realizing this potential would require a larger-scale analysis, we tested six common ML estimators from the scikit-learn library in python (Pedregosa et al. 2011) on the ADH mutations as a proof of concept. Predicting stabilizing mutations can be framed as either a classification (stabilizing or not), or a regression task (predicting ΔTm). We tested three classifiers—a linear kernel support vector classifier (SVC), K-nearest neighbors (KNC), and a random forest ensemble classifier. For regression, we applied support vector regression (SVR), ridge regression, and again random forest, due to their robustness and minimal tuning requirements (Parmar et al. 2019). Using default hyperparameters for all models, we performed repeated stratified k-fold cross validation using 4 splits and 20 repeats on all 140 experimentally verified ADH mutations. We set classification targets as 1 (stabilizing) or 0 (non-stabilizing), while regression models used ΔT_m_ values.

To assess whether the ML models could improve performance, we plotted the averaged weighted and unweighted ROC curves for all six estimators (Figure 9a-b), ranking by model probability and ΔT_m_ for classifier and regressor, respectively. While most models did not outperform the simple average of all metrics, the Random Forest models showed AUC improvements for the unweighted ROC curves in particular, indicating an improved performance in identifying stabilizing mutations (Figure 9c). Notably, although model prediction was not able to improve the weighted ROC AUC, the score remained consistently higher, indicating that highly stabilizing mutations were prioritized regardless of task and model (Figure 9c). We found that models did not generalize to other proteins (data not shown)—likely due to a too small dataset or indicating different main physical stabilization modes across proteins. However, the random forest performance boost suggests that once a sufficient number of mutations are identified, training a small ad hoc model can increase success rate, and demonstrates BoostMut’s utility for ML.

**Figure 9.**
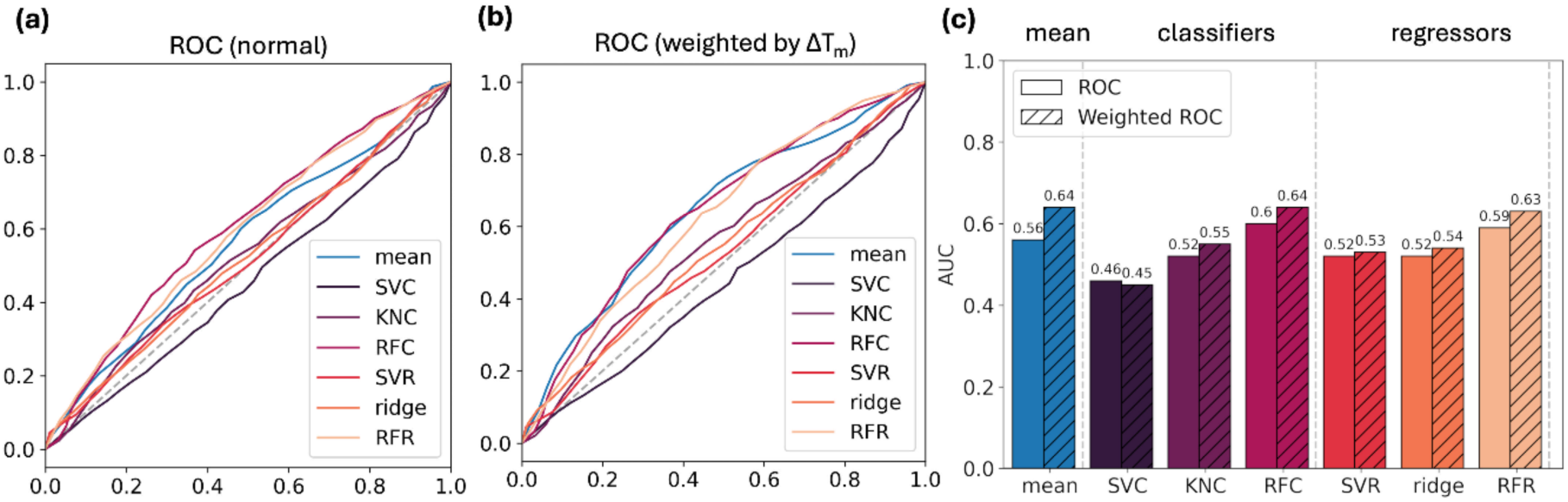
(a) ROC curves of the mean score and six BoostMut output-trained predictors averaged over the repeated stratified k-fold cross validation. (b) ROC curves weighted by ΔT_m and_ averaged over the repeated stratified k-fold cross validation. (c) AUC values corresponding to ROCs in (a) and (b).

## Discussion

We developed BoostMut to improve the identification of stabilizing mutations and assist in prioritizing individual mutations for experimental validation, complementing any primary predictor. This effort was motivated by the observation that while existing algorithms effectively filter out destabilizing mutations, their correlation with stabilizing effects is relatively weak (Khan and Vihinen 2010). BoostMut refines these pre-selected sets by providing a more reliable ranking. One of the main challenges was evaluating BoostMut as a secondary filter, since it operates on mutations already enriched for stability, complicating comparisons with other tools. We here used ROC curves to account for the imbalance between stabilizing and destabilizing mutations and propose weighted ROC analysis to better reflect stability improvements. While ROC curve weighting has precedent, it is commonly implemented as weights to classes or ROC curve sections rather than individual data points (Li and Fine 2010; Wang and Li 2012).

In our work, we evaluated BoostMut’s performance using four approaches: (1) We re-ranked 320 FRESCO-proposed mutations and found an improvement of ROC AUC on a mutation set where the high degree of pre-selection caused the primary predictor’s score to perform poorly. (2) We tested 333 mutations from the T2837 dataset, which included a more diverse set of mutations, and showed that BoostMut improved AUC when used alongside both the empirical force-field-employing FoldX or the machine learning predictor Stability Oracle. (3) We compared BoostMut to visual inspection on 1251 FRESCO mutations, where it selected significantly different mutations, but achieved a higher success rate (46%) among the overlapping mutations. (4) We experimentally validated 19 BoostMut-selected LEH mutations, of which 50% were stabilizing, demonstrating that BoostMut enhances selection even in cases where mutations were rejected manually. Across all tests, BoostMut consistently improved performance as a secondary filter.

Based on these results, we believe that BoostMut can replace visual inspections in thermostability engineering when time or expertise is limited, or augment it with objective guidance. Designed for broad compatibility, BoostMut remains independent of the primary stability predictor, allowing flexibility as methods, in particular ML-based, evolve. Beyond filtering, BoostMut may also enhance AI-driven thermostability predictions: its biophysical approach provides an orthogonal check on black-box ML models, helping identify prediction inconsistencies that would be physically obvious. Moreover, its machine-readable, standardized output can serve as feedback for further training, potentially aiding in the development of biophysics-aware ML algorithms. We take a step in this direction by demonstrating that a ML model tasked with optimizing metric weighting can be trained on BoostMut output.

Another strength of BoostMut is its implementation flexibility. While we show that longer MDs improve success rates, we find that the ranking converges relatively quickly, and leave this choice to the user, and even support static structure input. Similarly, users can customize metric priorities, facilitating system-specific considerations. For example, a salt bridge design campaign may warrant BoostMut use with other metrics disabled, while highly soluble proteins might justify deprioritizing hydrophobic exposure.

In summary, BoostMut provides a powerful tool for MD-supported secondary stability mutation filtering. Although MD simulations add a computational burden, the improvement in accuracy typically outweighs the additional cost and labor of experimentally assessing a larger fraction of false positive mutations. With ongoing developments in conformational ensemble generation, such short MD simulations could in the future be replaced by generative machine learning models with a much lower computational cost (Jing et al. 2024; Lewis et al. 2025). Besides ranking, BoostMut metrics may also serve as labels for large mutation datasets in the future training of improved ML models.

## Author Contributions

**Kerlen T. Korbeld:** Methodology; Investigation; software; data curation; visualization; writing – review and editing; writing – original draft. **Maximilian J.L.J. Fürst:** Conceptualization; investigation; funding acquisition; project administration; writing – review and editing; supervision; validation.

## Acknowledgement

We thank Dr. Hein J. Wijma for fruitful discussions and providing raw data of energy calculation and MD simulation data from previous FRESCO projects. M.J.L.J. Fürst gratefully acknowledges funding from the Netherlands Organization for Scientific Research NWO (VI.Veni.212.263). This work was supported by the European Union through COST Action CA21162 (Establishing a Pan-European Network on Computational Redesign of Enzymes) and used the Dutch national e-infrastructure with the support of the SURF Cooperative using grant no. EINF-4326. We also thank the Center for Information Technology of the University of Groningen for providing access to the Hábrók high performance computing cluster.

## Data Availability Statement

Source code, instructions for BoostMut, and all data and analysis used to generate the figures are available at https://github.com/kt-korbeld/BoostMut. The MD trajectories associated with the study are available upon request.

## Materials and Methods

BoostMut is implemented as a python package with a command line program called boostmut_run. All analyses are performed on 3 selections: the whole protein, the local surrounding of the mutation, and just the interactions with the mutation itself. For the local surrounding, we use a default cutoff radius of 8 Å, which has been suggested as an optimal context to assess effects on stability (Gromiha et al. 2000). A second command line interface called boostmut_process provides tools to process the raw output, such as adding primary predictors as metrics, scaling the values by standard deviation, or outputting the data into a human readable excel format. An online tool implemented as an R shiny app for the visualization of the metrics and data is freely available at https://fuerstlab.shinyapps.io/BoostMutUI/.

### Hydrogen Bonding

The analysis of both the H bond energy and unsaturated H bonds are calculated and averaged over each frame in the trajectory. H bonds are identified using the hbond_analysis function in the MDAnalysis package. The donor-hydrogen-acceptor angle and donor-acceptor distance cutoffs used to identify H bonds are set to be more lenient than the default settings (an angle cutoff of 100° instead of the default 150° and distance cutoff of 3.5 Å instead of the default 3 Å respectively) as later steps filter out unrealistic H bonds using an approximation of their bonding energy.

The bonding energy is approximated based on the angles and distances of the H bond, allowing BoostMut to work without charge information. The energy function is inspired by the ListHBo command in YASARA, which we further modified. The formula scales down an ideal interaction energy of 25 kJ/mol based on deviations in the distance (Scale_HA_) and angle (Scale_DHA_) away from the ideal conformation (Hubbard and Kamran Haider 2010). Energy is further scaled down if any atom covalently bound to the acceptor is close enough to the hydrogen to hinder the H bond (Scale_HAX_). The formula as provided by YASARA is then as follows:

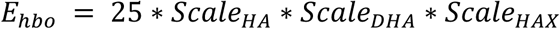

The formula for each of the scaling factors are saturated ramp functions between 0-1, The ramp function of Scale_HAX_ depends on whether the covalent atom X is a hydrogen or a heavy atom. Since the latter causes more steric hinderance, stricter criteria are applied. If there are multiple covalent atoms attached to the acceptor, the lowest scaling factor is used.

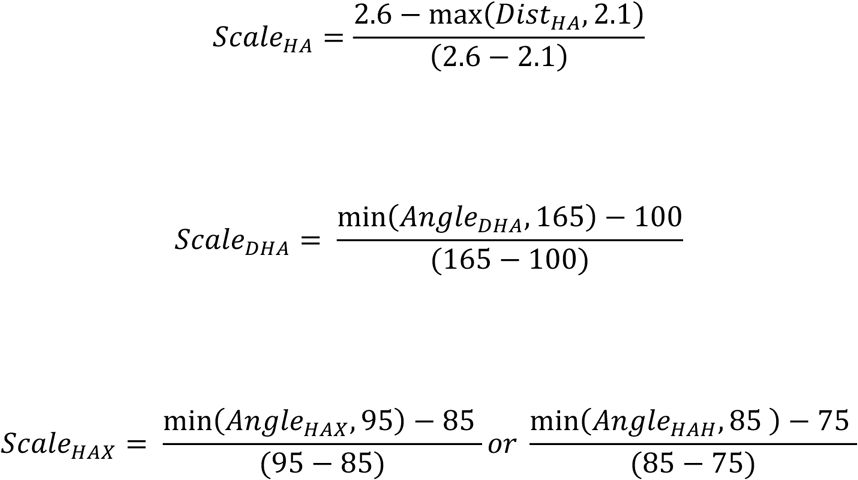

For bifurcated H bonds the second bond is scaled down to a maximum of 15 kJ/mol (Feldblum and Arkin 2014). The formula was applied to all H bonds identified by MDAnalysis, and only H bonds with an energy higher than 6.25 kJ/mol are selected.

First order Water-mediated H bonds or water bridges are included in the H bond energy (Petukhov et al. 1999). Water bridges are identified by first selecting all waters involved in two or more H bonds with the protein and then selecting all protein-water H bonds containing these waters. The bond energy of a water bridge is estimated by summing the energies of all H bonds involved in the water bridge and subtracting an entropic penalty of 32.2 kJ/mol for removing a water from solution (Petukhov et al. 1999). To find the number of unsatisfied hydrogen bonds, the set of all atoms engaged in a H bond are compared to the set of all potential H bonds partners (defined as all Oxygen atoms, and all Nitrogen atoms not bonded to any Hydrogens). Whenever an atom was found in the set of potential H bonds partners, but not in the set of atoms engaged in a hydrogen bond, the atom was counted as an unsatisfied hydrogen bond.

### Protein Flexibility

Both the RMSF of the backbone and the flexibility score of the sidechain are calculated. The flexibility of the backbone is measured by calculating the RMSF of the Ca atoms in the backbone using the MDAnalysis package. For the RMSF values of the sidechains, a local alignment of +-1 amino acids around the residue is performed for each residue so that the RMSF score is not dominated by fluctuations in the backbone, and the RMSF for the sidechain atoms excluding the backbone is calculated.

For the sidechain score, benchmark distributions were categorized based on amino acid type and whether the residue was buried, partially exposed, or fully exposed, resulting in a total of 20 * 3 = 60 different benchmark distributions (Figure S3). The three categories were defined using the fraction of solvent exposed surface (SASA), calculated with freeSASA (Mitternacht 2016; Shrake and Rupley 1973). The residue is considered buried if 0% of the residue is solvent exposed, partially exposed if between 0-20% is solvent exposed, and fully exposed if the residue is >20% is solvent exposed (Chen and Zhou 2005; Kumar and Nussinov 1999; Miller et al. 1987). The percentage of exposure is defined as the SASA value divided by the average SASA of the single amino acid in solution reported in literature.

To create the benchmark curves, A Gaussian KDE was fitted to the distributions for each category obtained from the benchmark data set (described below), with the bandwidth set using Silverman’s rule of thumb. For each of the 60 categories the maximum value was scaled to 1, allowing for a mapping from the sidechain RMSF to a sidechain score between 0 and 1. In order to guarantee the score improves monotonically when the sidechain RMSF is lowered, each value was set to the maximum between either its own value or the highest score among all RMSF values higher than the current one (i.e., “filling in” the local minima). The resulting curves are saved as a series of x and y values in .csv format for each of the categories.

### Hydrophobic Surface Exposure

The degree of hydrophobic surface exposure is determined by calculating SASA with FreeSASA and averaged over the trajectory. Hydrophobic moieties were approximated as all atoms engaged in a C-C or C-H bond. A set of curves with distributions of hydrophobic surface exposure per amino acid were created from the benchmark dataset, resulting in 20 categories. The benchmark curves were processed identically to the RMSF benchmark curves: A Gaussian KDE was fitted to the distributions for each category obtained from the test set, with the bandwidth set using Silverman’s rule of thumb. The maximum value was scaled to 1, and the local minima were flattened to guarantee a monotonic decrease.

### Other Metrics

Besides H bonds, flexibility and surface exposure, four other structural metrics are recorded which could disrupt protein stability. All *α*-helices are identified on the first frame of the simulation using pyDSSP (Kabsch and Sander 1983). To find helix capping motifs, all possible motifs are checked within 5 amino acids around the start or end of the helix to account for fluctuations in the assignment of the secondary structure (Yang et al. 2016). To track helix propensity, we take the scale reported in Pace et al. and track its change for any residues inside an *α*-helix and 5 residues away from the start or end of the helix (Pace and Scholtz 1998). To verify that no cysteine-cysteine disulfide bonds have been disrupted by any of the mutations, we count the number bonds between cysteine residues before and after mutation. The number of salt bridges is counted by taking all cases where an atom capable of being an electron donor is less than 4 Å away from an atom capable of being an electron acceptor (Kumar and Nussinov 1999). Only the number of salt bridges is averaged over each frame of the trajectory and reported for all three selections.

### High Throughput Molecular Dynamics

High throughput molecular dynamics were performed using the YASARA software package, using the same protocol previously reported for FRESCO (Wijma et al. 2018). This pipeline involves the following steps: addition and optimization of hydrogens using the YASARA software, solvation and neutralization of the simulation cell with 0.5 M NaCl, followed by a steepest descent minimization. The system is then brought up to a temperature of 298 K in a 20 ps equilibration run, followed by a 50 ps production run. A snapshot was taken every 5 ps and saved in the YASARA-specific “.sim” format. All simulated steps (minimization and MD) were done using the Amber derived YAMBER3 forcefield, with long range interactions calculated using PME.

For further analysis by BoostMut, the .sim files were converted to a format recognized by MDAnalysis. The .sim files were converted into a .xtc file using the md_convert macro provided by YASARA. Besides the trajectory, a topology file containing least atom and bond information should be provided for the automated inspection, which can either be done by using GROMACS to generate a .trp from a given .pdb, by saving bond information into the pdb, or by using YASARA to save the bond information separately, for which some helper functions have been provided.

For the three proteins with more than 100 mutations (1BZ6, 2LZM, 1LZ1) reported in the T2837 test set, the structures for all mutations were generated using FoldX and run using the high-throughput MDs. For LEH, the original MD trajectories were preserved. For ADH and HMFO, MD simulations were re-run using the same initial structures and simulation parameters as the original FRESCO campaign.

### Benchmark data set

A benchmark set of proteins was obtained from the PDB. In order to obtain a well behaved, representative proteins all proteins were selected with 1) A refinement resolution of <= 1 Å, 2) A length of 100 amino acids or larger, and 3) no non-proteogenic entities present: (total number of non-polymer entities = 0, polymer entity type = protein). This resulted in 77 proteins with a length distribution between 100 to 501 amino acids.

The same pipeline for high-throughput Molecular Dynamics as described above were run on each of the structures for 1000 ps. To get the data for the RMSF benchmark curves, each residue in each of the proteins in the test set was categorized based on surface exposure and amino acid type, and its RMSF recorded. For hydrophobic exposure, each residue was categorized based on amino acid type and its hydrophobic surface exposure was recorded.

Benchmarks were generated for simulation times between 50-1000 ps in 50 ps intervals by slicing the 1000 ps trajectory to the desired length.

### Experimental Characterization of new mutations

A synthetic gene encoding LEH was ordered from Integrated DNA Technologies after BSAI sites were added to the 5’- and 3’-termini. The gene encoding LEH was cloned into a pBAD His vector using golden gate cloning (Engler and Marillonnet 2014). To further facilitate automation, we created a streamlined python implementation of AAscan a software package used to design partially overlapping mutagenesis primers previously used in FRESCO and other high-throughput protein engineering campaigns (Wijma et al. 2018). The python implementation (https://github.com/kt-korbeld/pyAAscan) (Sun et al. 2013) is based on the original AAscan code (https://github.com/dbv123w/AAScan). Primers for the mutations were designed to be overlapping and each mutation was introduced using the QuikChange protocol utilizing PfuUltra Hotstart PCR Master Mix. The product was transformed into NEB10β, and all mutations were verified through direct colony sequencing using a ColonySeq plate provided by Eurofins. Each mutation was expressed by inoculating an overnight culture of the transformed NEB10β colonies into 50 mL of TB containing 50 μg/mL ampicillin. Once the cultures reached an OD_600_ of 0.6-1.0 they were induced with 0.02% L-arabinose and incubated for 16 h at 30 °C. The cells were harvested by centrifugation 15,000 × g for 1 h, 4 ℃ and stored at −20 ℃ until needed.

During purification, the buffers used were those reported in the most recent engineering campaign on LEH (Wijma et al. 2015). To purify the protein, the pellets were resuspended in lysis buffer (50mM Hepes, 500mM NaCl, pH 8) until they reached an OD_600_ of 12, lysed using sonication (5min total time, 5s on, 10s off) and the cell debris removed by centrifugation (15000G, 1hr, 4 ℃). The cell free extract was loaded onto an equilibrated Ni-Sepharose column and left to incubate for 1 hour at 4 ℃. The column was washed with washing buffer (50mM hepes, 500 mM NaCl, 20 mM imidasol, pH 8) for 10 Column Volumes (CV), after which the protein was eluted with 5 CVs of elution buffer (50mM hepes, 500mM NaCl, 300mM imidasol, pH8). The obtained protein was desalted using a desalting column, obtaining the final protein in a desalted buffer (50mM Hepes, pH8).

To measure the melting temperature, 20 μL each of the isolated protein variants (with a concentration between 0.5-2.5 mg/ml) was loaded into transparent PCR tubes together with 5 μL of 100-fold diluted Sypro Orange (Life Technologies, CA, USA) after which the plate was sealed with transparent foil. The fluorescence (excitation at 490 nm and emission at 575 nm) was monitored while the sample was heated from 20 to 99 ℃ at 1.1 ℃/min in a RT-PCR machine (CFX96 Touch Real-Time PCR, BioRad, signal setting: FRET). The melting temperature was obtained by taking the maximum of the first derivative of the fluorescence curve over time.

### ROC AUC analysis

Normal ROC AUC is plotted using the sklearn.metrics.roc_curve function. For the weighted ROC AUC, we implement a custom version of the ROC function that weighs each change in the true positive rate by the relative contribution to the total increase in ΔT_m_. As we are not interested in the degree are mutation is destabilizing, the true negatives are not scaled.

To identify the best and worst-case scenarios among the remaining untested mutations, we calculate the upper and lower bounds of the ROC AUC. We ignore all permutations where the best and worst-case scenarios do not have all true positives sorted as close as possible to either the highest or lowest rank respectively. We therefore iteratively change each untested mutation to be stabilizing from highest to worst ranked and from worst to highest ranked, calculate the AUC for each scenario.

We also plot the ROC AUC as a function of an increasing fraction of stabilizing mutations within the untested mutations. Here we are no longer interested in a particular best- or worst-case scenario for the ROC AUC, but calculate the average effect of a given fraction of positive hits, independent of its exact sorting. To calculate this for each fraction, the desired number of positive hits are randomly introduced among the unknown mutations and the new ROC AUC is calculated. This is repeated 1000 times to average out the effect of any particular sorting.

### Machine learning

The scikit.learn implementation of each classifier and regressor was used with default hyperparameters. The support vector machines were implemented with sklearn.svm.SVC, and sklearn.svm.SVR using a linear kernel. The K-nearest Neighbors classifier was implemented using sklearn.neighbors.KNeighborsClassifier, the ridge regressor using sklearn.linear_model.Ridge. The Random Forest approaches were implemented using sklearn.ensemble.RandomForestClassifier and sklearn.ensemble.RandomForestRegressor. For both classifiers and regressors, the scaled BoostMut metrics were provided as features. For classifier models, classes of stabilizing or destabilizing mutations were provided as labels. For the regressor models, the ΔT_m_ values were provided as labels. The 140 experimentally verified mutations for ADH were used as data. The models were cross-validated using repeated stratified k-fold cross validation, using 4 splits and 20 repeats, and the averaged ROC curves and AUC values are reported. For the classifiers, the model probability was used to generate the ROC curves.

### Other analyses and visualization

All figures were produced using the matplotlib and seaborn python libraries. Venn diagrams were generated using matplotlib-venn. Protein structures were visualized using YASARA or ChimeraX. The secondary structure representation in figure 6 was generated using the secstructartist python package. Predictions using DDGun(Montanucci et al. 2022), DDMut (Zhou et al. 2023), DUET (Pires et al. 2014), Mupro (Cheng et al. 2006), mCSM, and PremPS (Chen et al. 2021) were generated using the Benchstab command line interface (Velecký et al. 2024). Predictions for FoldX were calculated using protocols established for the FRESCO pipeline (Wijma et al. 2018). Predictions for Pythia (Sun et al. 2023) were generated using the provided colab notebook at https://colab.research.google.com/gist/JinyuanSun/83ff4323ff751dc665f96381a02df18a/colabpythia.ipynb#scrollTo=iJ58Cx4z2lof

