## Supplementary Figures S1-S3 for "Enriching stabilizing mutations through automated analysis of molecular dynamics simulations using BoostMut"

### Figure S1

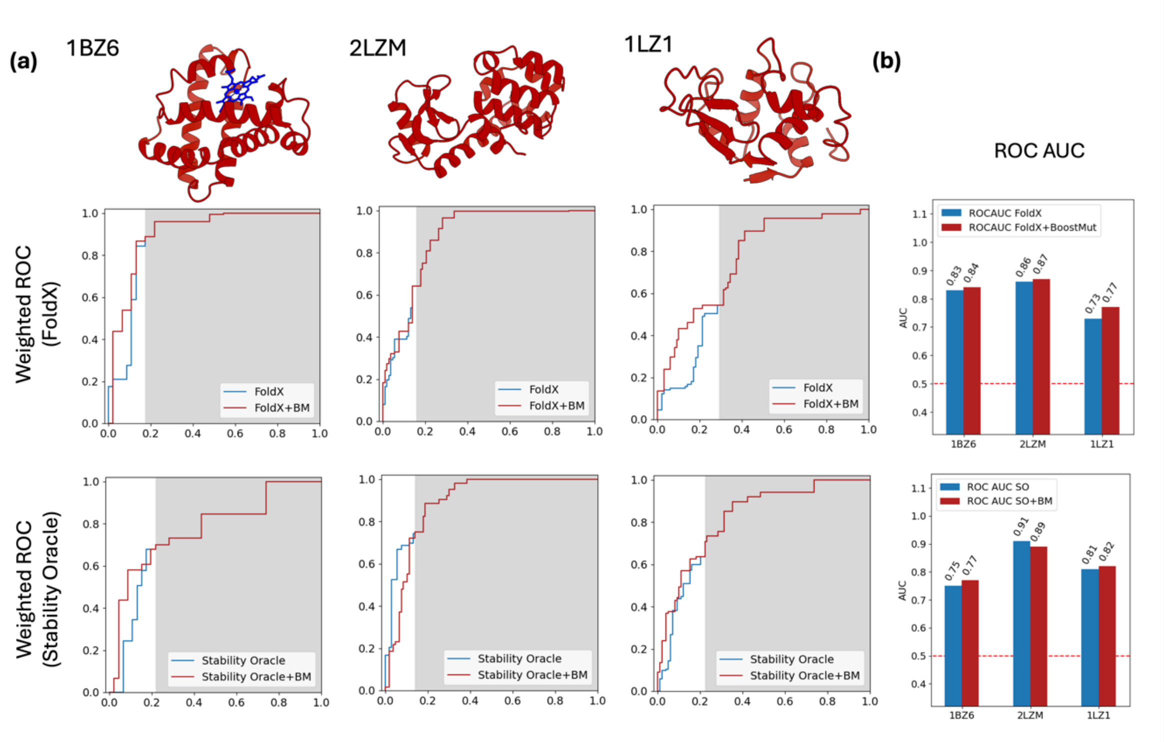

*Figure S1 (a) Weighted ROC curves for three proteins from the T2837 test set, using either FoldX (top row) or Stability Oracle (bottom row) as the primary predictor. All mutations with a predicted ΔΔG < 0 were scored with the automated assessment (the area shown in white), all mutations with a predicted ΔΔG > 0 were scored by the primary predictor (the area shown in grey). (b) For all cases apart from Stability Oracle when used on 2LZM, the addition of the automated assessment resulted in a small increase in the weighted ROC AUC.*

### Figure S2

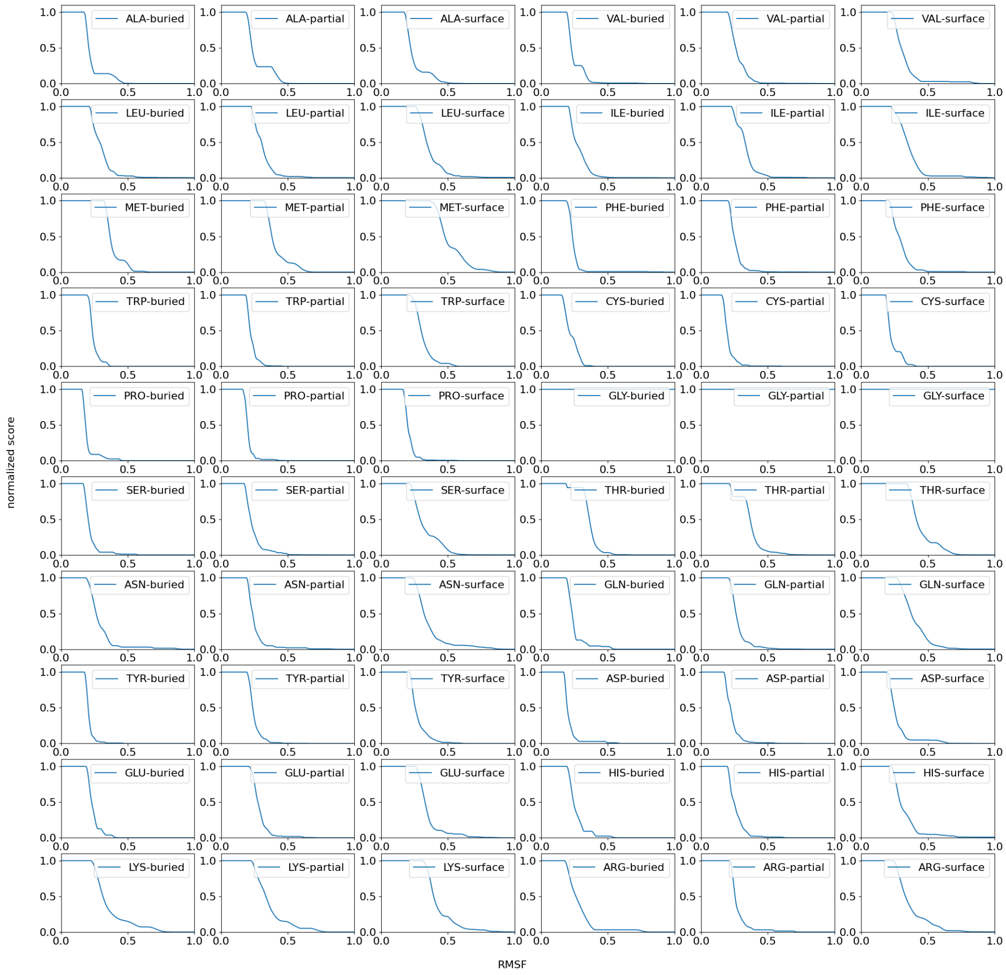

*Figure S2 Benchmark curves for expected sidechain flexibility (RMSF) for 60 categories, resulting from classification of each amino acid into “exposed” (>20% relative SASA), “partial” (>0% and <20% relative SASA), and “buried” (0% relative SASA). Distributions of RMSF for each of the categories are obtained from the benchmark data, fitted with a Gaussian kernel, scaled to a maximum value of 1, and made to decrease monotonically.*

### Figure S3

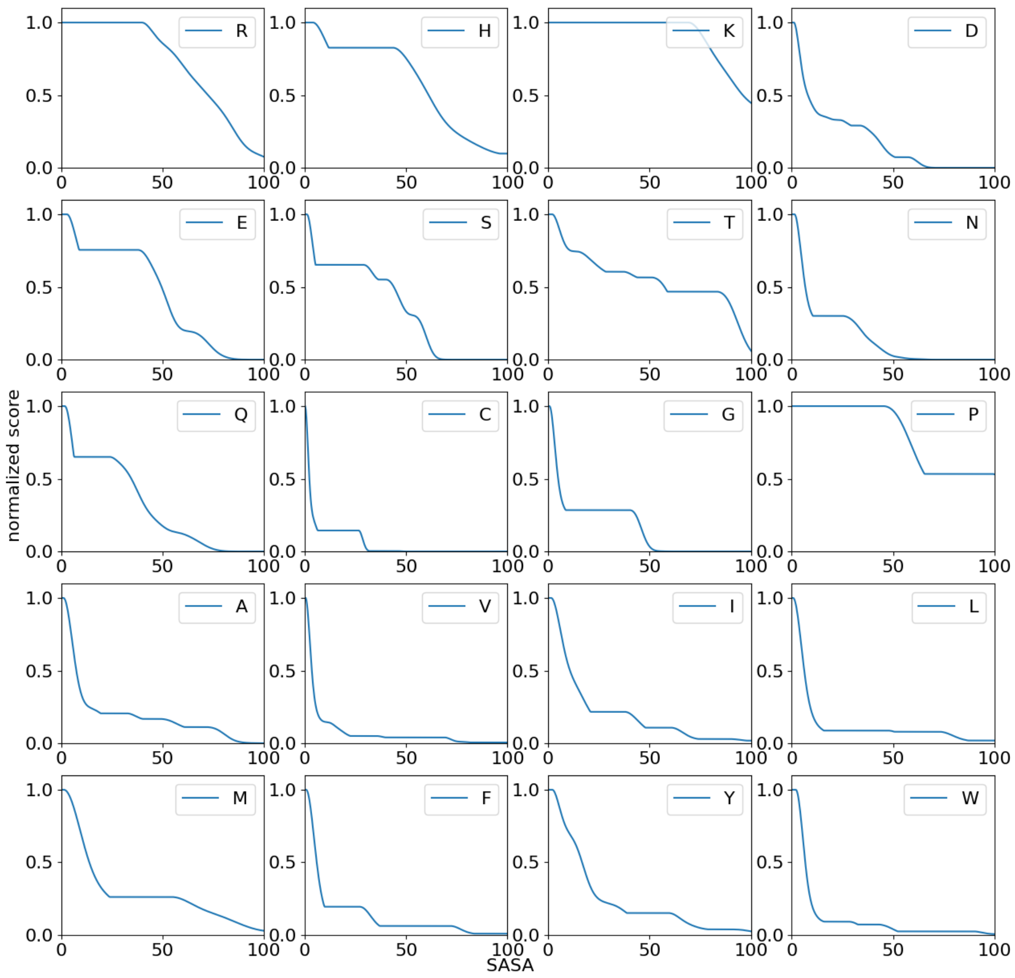

*Figure S3 Benchmark curves for expected hydrophobic exposure (SASA) for each of the amino acids. Distributions of SASA for each of the categories are obtained from the benchmark data, fitted with a Gaussian kernel, scaled to a maximum value of 1, and made to decrease monotonically.*
